# Protein composition of the occlusion bodies of *Epinotia aporema* granulovirus

**DOI:** 10.1101/465021

**Authors:** Tomás Masson, María Laura Fabre, María Leticia Ferrelli, Matías Luis Pidre, Víctor Romanowski

## Abstract

Within family *Baculoviridae*, members of the *Betabaculovirus* genus are employed as biocontrol agents against lepidopteran pests, either alone or in combination with selected members of the *Alphabaculovirus* genus. *Epinotia aporema* granulovirus (EpapGV) is a fast killing betabaculovirus that infects the bean shoot borer (*E. aporema*) and is a promising biopesticide. Because occlusion bodies (OBs) play a key role in baculovirus horizontal transmission, we investigated the composition of EpapGV OBs. Using mass spectrometry-based proteomics we could identify 56 proteins that are included in the OBs during the final stages of larval infection. Our data provides experimental validation of several annotated hypothetical coding sequences. Proteogenomic mapping against genomic sequence detected a previously unannotated ac110-like core gene and a putative translation fusion product of ORFs *epap48* and *epap49.* Comparative studies of the proteomes available for the family *Baculoviridae* highlight the conservation of core gene products as parts of the occluded virion. Two proteins specific for betabaculoviruses (Epap48 and Epap95) are incorporated into OBs. Moreover, quantification based on emPAI values showed that Epap95 is one of the most abundant components of EpapGV OBs.

## Introduction

The family *Baculoviridae* comprises a diverse group of large double stranded DNA viruses that infect larvae of the insect orders *Lepidoptera*, *Hymenoptera* and *Diptera* [1]. Baculovirus have a rod shaped, enveloped virion with a circular genome ranging from 80 to 180 kbp [2]. Virions are found in the environment embedded in a proteinaceous matrix that forms occlusion bodies (OBs), a phenotype that is resistant to desiccation and UV radiation. OBs on leaves that are consumed by foraging larvae reach the midgut and, after being dissolved at high pH, release the occlusion derived viruses (ODVs), which initiate infection of the epithelial cells. These infected cells produce budded viruses (BVs) that disseminate the infection systemically [3]. Based on OBs morphology, baculoviruses were first classified in two groups: Nucleopolyhedrovirus (NPV) and Granulovirus (GV) [1]. Later, they were taxonomically divided into four genera: *Alphabaculovirus* (lepidopteran-specific NPV), *Betabaculovirus* (lepidopteran-specific GV), *Gammabaculovirus* (hymenopteran-specific NPV) and *Deltabaculovirus* (dipteran-specific NPV) [1].

Among different entomopathogenic viruses, the baculoviruses have received most of the attention due to their narrow host range which makes them safe pesticides. The majority of commercial products are based on virus isolates that belong to the genera *Alphabaculovirus* and *Betabaculovirus* [4]. The bean shoot borer (*Epinotia aporema*) is an oligophagous pest that attacks soybean crops [5]. A poliorganotropic fast killing betabaculovirus for this species, *Epinotia aporema* granulovirus (EpapGV), has been discovered and sequenced by our group [6, 7]. In order to improve our understanding of the infectious process we set out to analyze the protein content of EpapGV OB using a proteomic approach.

Mass spectrometry-based (MS) proteomics represents a powerful technique to interrogate the structural landscape of viral particles [8]. In addition to direct protein identification, spectral data derived from proteomic experiments can be used to identify novel features within genomic and transcriptomic datasets. This proteogenomic methodology is independent of reference annotation, thus providing excellent means for the refinement of gene models and the discovery of novel protein coding sequences [9].

Virion proteomics has been applied to study several DNA virus families (*Ascoviridae* [10], *Herpesviridae* [11], *Iridoviridae* [12], *Nudiviridae* [13] and *Poxviridae* [14]). In relation with the present study, eight ODV proteomes of baculoviruses have been analyzed (AcMNPV [15], AgMNPV [16], ChchNPV [17], HearNPV [18], MabrNPV [19], ClanGV [20], PiraGV [21] and CuniNPV [22]). These datasets point at a complex virion comprising a large number of proteins involved in virion morphogenesis, OBs formation and infection of insect midgut epithelial cells.

In this work we examined the protein content of EpapGV OBs using MS-based shotgun proteomics in order to describe the composition of this virion phenotype. A total of 56 viral proteins from EpapGV OBs were identified, the majority of which are conserved components among the members of the family *Baculoviridae* and few are betabaculovirus-specific proteins. Comparative proteomics of baculovirus showed a set of core gene products present in the majority of proteomes analyzed.

## Materials and Methods

### Larvae and virus

Larvae of the bean shoot borer (*Epinotia aporema*), were collected from field in the experimental station of the Instituto Nacional de Tecnología Agropecuaria (INTA) and reared in our laboratory with an artificial diet and controlled light cycle (16 hours of light). The strain used in this study was EpapGV (Refseq ID NC_018875), collected in Oliveros (Santa Fe, Argentina) [6].

### Occlusion bodies (OBs) production and purification

Fourth instar *E. aporema* larvae were infected *per os* using artificial diet contaminated with a solution containing EpapGV OBs. Dying larvae with signs of infection were stored and processed as described previously [7]. Briefly, infected larvae were stored in distilled water and later homogenized in a Dounce homogenizer. The resulting solution was filtered through three layers of cheesecloth to eliminate insoluble insect debris. This extract was clarified by three steps of centrifugation at 10000 x g for 10 minutes followed by a wash with 0.05% v/v SDS solution. Clarified solution was subjected to ultracentrifugation in a continuous 30-60% w/w sucrose gradient (50000 x g, one hour, 4°C, Beckman SW 41 Ti rotor). The whitish/opalescent band corresponding to OBs was collected, diluted 10-fold in distilled water and pelleted by centrifugation at 14000 x g for 10 minutes. The final pellet was resuspended in distilled water and stored frozen at −20°C. Two biological independent samples were processed. Total protein mass in the sample was quantified using the Bradford assay [23].

### Mass spectrometry analysis

Protein digestion and analysis were performed at the Proteomics Core Facility CEQUIBIEM, at the University of Buenos Aires/CONICET (National Research Council) as follows: protein samples were reduced with 10 mM dithiothreitol in 50 mM ammonium bicarbonate pH 8 (45 min, 56°C) and carbamidomethylated with 20 mM iodoacetamide in the same solvent (40 min, room temperature, in darkness). This protein solution was precipitated with 0.2 volumes of 100% w/v trichloroacetic acid (Sigma) at −20 °C for at least two hours and centrifuged at 12000 x g for 10 min (4°C). The pellet was washed twice with ice-cold acetone and dried at room temperature. Proteins were resuspended in 50 mM ammonium bicarbonate pH 8 and digested with trypsin (Promega V5111). The resulting peptides were desalted with ZipTip C18 columns (Millipore).

The digests were analyzed by nanoLC-MS/MS in a Thermo Scientific Q Exactive Mass Spectrometer coupled to a nanoHPLC EASY-nLC 1000 (Thermo Scientific). For the LC-MS/MS analysis, approximately 1 μg of peptides was loaded onto the column and eluted for 120 minutes using a reverse phase column (C18, 2 μm x 10 nm particle size, 50 μm x 150 mm) Easy-Spray Column PepMap RSLC (P/N ES801) suitable for separating complex mixtures of peptides with a high degree of resolution. The flow rate used for the nano-column was 300 nL min^-1^ and the solvent range from 7% B (5 min) to 35% B (120 min). Solvent A was 0.1% formic acid in water whereas B was 0.1% formic acid in acetonitrile. The injection volume was 2 μL. The MS equipment has a high collision dissociation cell (HCD) for fragmentation and an Orbitrap analyzer (Thermo Scientific, Q-Exactive). A voltage of 3.5 kV was used for Electro Spray Ionization (Thermo Scientific, EASY-SPRAY).

XCalibur 3.0.63 (Thermo Scientific) software was used for data acquisition and equipment configuration that allows peptide identification at the same time of their chromatographic separation. Full-scan mass spectra were acquired in the Orbitrap analyzer. The scanned mass range was 400-1800 m/z, at a resolution of 70000 at 400 m/z and the 12 most intense ions in each cycle, were sequentially isolated, fragmented by HCD and measured in the Orbitrap analyzer. Peptides with a charge of +1 or with unassigned charge state were excluded from fragmentation for MS2.

### Analysis of MS data

Q Exactive raw data was processed using Proteome Discoverer^™^ software (version 2.1.1.21, Thermo Scientific) and searched against EpapGV protein database downloaded from NCBI (accession number NC_018875, National Center for Biotechnology Information; www.ncbi.nlm.nih.gov) digested with trypsin with a maximum of one missed cleavage per peptide. Proteome Discoverer^™^ searches were performed with a precursor mass tolerance of 10 ppm and product ion tolerance of 0.05 Da. Static modifications were set to carbamidomethylation of Cys, and dynamic modifications were set to oxidation of Met and N-terminal acetylation. Protein hits were filtered for high confidence peptide matches with a maximum protein and peptide false discovery rate of 1% calculated using a reverse database strategy. The exponentially modified protein abundance index (emPAI) was calculated automatically by Proteome Discoverer^™^ software and used to estimate the relative abundance of identified proteins within the sample.

### Non annotated peptides search

The complete genome sequence of EpapGV was translated *in silico* in all six frames using the Mascot search software. Spectral data was searched and all peptides hits were filtered to discard matches in previously annotated ORFs. The remaining peptides were mapped to the corresponding genomic position. Search for putative unannotated ORFs was done extending peptide hits until a stop codon was found at C-terminus, and a start or stop codon for the N-terminus. Homologous sequences were searched using the TBLASTN tool against all baculovirus genomes.

### Orthologs clustering

A database comprising all the ODV proteins detected in baculoviruses was generated using previous proteomic data sets [15-22]. The software BLASTP [24] and HHMER [25] were used to identify groups of orthologous proteins (orthogroups) among different proteomes by reciprocal best hits.

## Results

### Structural components of the EpapGV OB

We determined the composition of purified EpapGV OBs employing a shotgun proteomic approach. The peptide mixture was separated by liquid chromatography and analyzed with tandem mass spectrometry (LC-MS/MS). This approach was used to avoid protein loss associated with SDS-PAGE gel extraction. We detected 56 proteins in our purified EpapGV OBs samples. Genes encoding these proteins comprise 43.93% of EpapGV total number of annotated ORFs (Fig 1), showing that a large part of the viral genome codes for structural proteins. A total of 10 proteins (Epap10, Epap62, Epap71, Epap123, LEF6, P6.9, Hel-1, P18, DNA Polymerase and DNA Ligase) were detected with only a single peptide, which might be related to low molar proportions of these polypeptides in the sample (Table 1). As additional evidence for the identification of these proteins, we checked the presence of several ion products belonging to the theoretical *b* and *y* spectral ions series for these peptides. The full list of proteins is shown in Table 1. In our samples we were unable to detect PIF3 and desmoplakin, two core gene products which have been confirmed in other virions by MS and western blot [15]. This could be attributed to proteolytic degradation, low protein level or deficient ionization of these proteins in our samples.

**Fig. 1.**
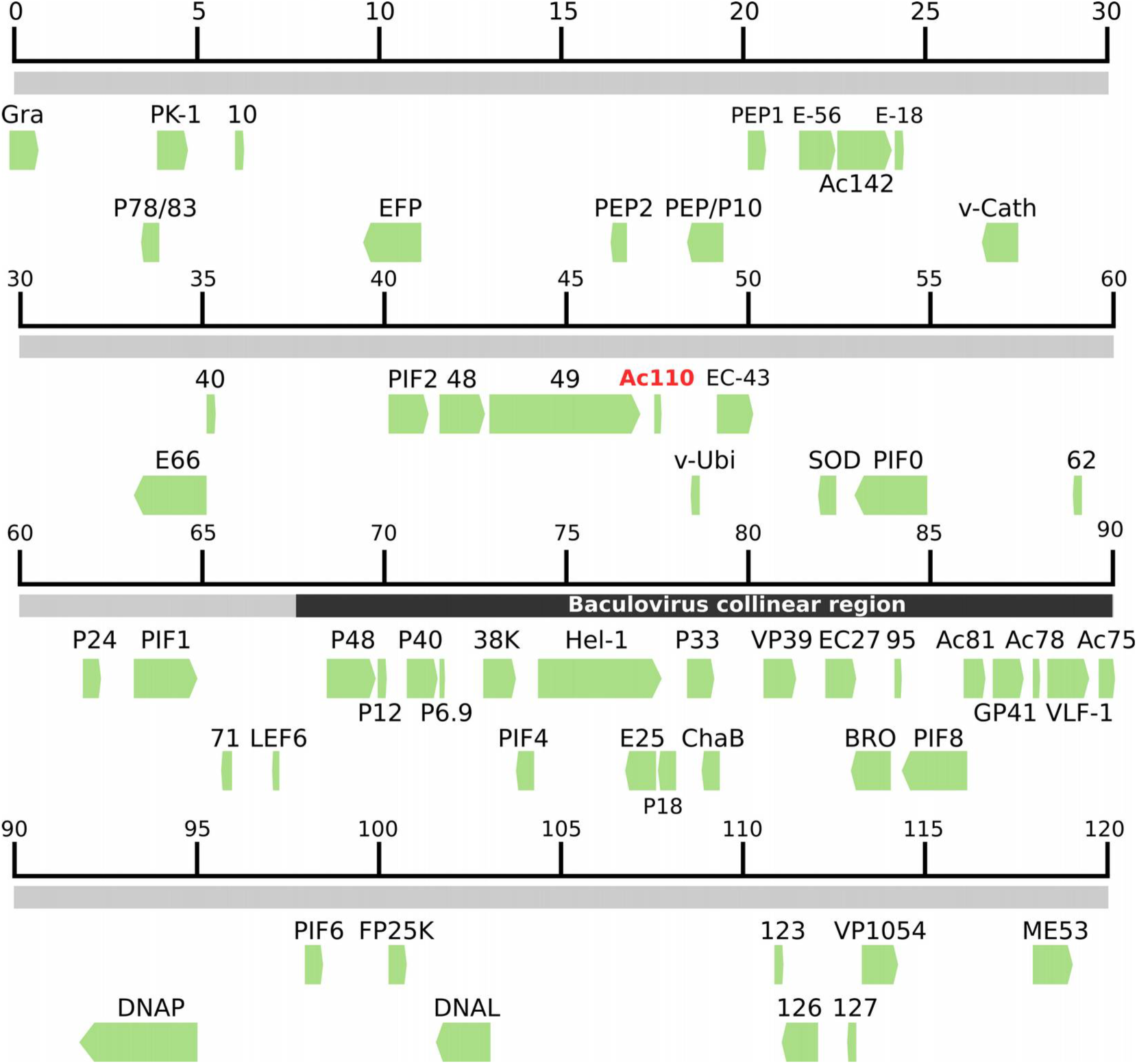
Genomic localization of ORFs coding for proteins found in EpapGV OBs. Proteins identified from spectral data (green arrows) were mapped to their respective genomic coordinate (grey line). The novel identified Ac110-like peptide is highlighted in red script. The baculovirus collinearity region is shown in black.

**Table 1.** Proteins detected in EpapGV OBs

| ORF | Protein | NCBI Protein Id. | Size (aa) | % Coverage | # Peptides | emPAI | % emPAI |
| --- | --- | --- | --- | --- | --- | --- | --- |
| 1 | Granulin | YP_006908509.1 | 248 | 60,48 | 25 | 1995261 | 857918 |
| 5 | P78/83 | YP_006908513.1 | 137 | 16,06 | 2 | 0,33 | 0,14 |
| 6 | PK1 | YP_006908514.1 | 276 | 30,43 | 9 | 3,44 | 1,48 |
| 10 | Epap10 | YP_006908518.1 | 90 | 23,33 | 1 | 0,33 | 0,14 |
| 14 | EFP | YP_006908522.1 | 541 | 5,18 | 1 | 0,08 | 0,03 |
| 21 | PEP2 | YP_006908529.1 | 142 | 45,07 | 5 | 4,62 | 1,99 |
| 22 | PEP/P10 | YP_006908530.1 | 308 | 48,70 | 8 | 9 | 3,87 |
| 25 | PEP1 | YP_006908533.1 | 178 | 37,64 | 6 | 5,31 | 2,28 |
| 27 | **PIF5** | YP_006908535.1 | 354 | 31,07 | 7 | 5,58 | 2,40 |
| 28 | **Ac142** | YP_006908536.1 | 457 | 44,86 | 18 | 4,84 | 2,08 |
| 29 | **ODV-E18** | YP_006908537.1 | 88 | 45,45 | 6 | 176,83 | 76,03 |
| 31 | v-Cath | YP_006908539.1 | 329 | 5,77 | 2 | 0,29 | 0,12 |
| 39 | ODV-E66 | YP_006908547.1 | 654 | 33,64 | 14 | 5,31 | 2,28 |
| 40 | Epap40 | YP_006908548.1 | 102 | 75,49 | 6 | 16,78 | 7,22 |
| 47 | **PIF2** | YP_006908555.1 | 374 | 20,05 | 7 | 0,69 | 0,30 |
| 48 | Epap48 | YP_006908556.1 | 446 | 37,67 | 13 | 2,54 | 1,09 |
| 49 | Epap49 | YP_006908557.1 | 1465 | 29,83 | 35 | 2,88 | 1,24 |
| 52 | v-Ubi | YP_006908560.1 | 93 | 33,33 | 3 | 1,37 | 0,59 |
| 53 | **ODV-EC43** | YP_006908561.1 | 348 | 47,41 | 12 | 6,20 | 2,67 |
| 58 | SOD | YP_006908566.1 | 183 | 55,19 | 9 | 24,12 | 10,37 |
| 59 | **PIF0** | YP_006908567.1 | 653 | 11,02 | 5 | 0,47 | 0,20 |
| 62 | Epap62 | YP_006908570.1 | 106 | 10,38 | 1 | 0,39 | 0,17 |
| 66 | P24 | YP_006908574.1 | 165 | 27,88 | 3 | 0,87 | 0,37 |
| 69 | **PIF1** | YP_006908577.1 | 538 | 7,99 | 3 | 0,52 | 0,22 |
| 71 | Epap71 | YP_006908579.1 | 104 | 6,73 | 1 | 0,47 | 0,20 |
| 74 | LEF6 | YP_006908582.1 | 82 | 12,19 | 1 | 0,47 | 0,20 |
| 78 | **P48/45** | YP_006908586.1 | 380 | 6,58 | 2 | 0,10 | 0,04 |
| 79 | P12 | YP_006908587.1 | 115 | 38,26 | 3 | 4,62 | 1,99 |
| 80 | **P40** | YP_006908588.1 | 373 | 25,20 | 9 | 1,68 | 0,72 |
| 81 | **P6.9** | YP_006908589.1 | 56 | 14,29 | 1 | 2,16 | 0,93 |
| 83 | **38K** | YP_006908591.1 | 295 | 26,44 | 5 | 1,45 | 0,62 |
| 84 | **PIF4** | YP_006908592.1 | 162 | 20,99 | 3 | 0,67 | 0,29 |
| 85 | **Hel-1** | YP_006908593.1 | 1085 | 1,29 | 1 | 0 | 0,00 |
| 86 | **ODV-E25** | YP_006908594.1 | 213 | 43,19 | 7 | 9 | 3,87 |
| 87 | **P18** | YP_006908595.1 | 158 | 5,06 | 1 | 0 | 0,00 |
| 88 | **P33** | YP_006908596.1 | 254 | 36,61 | 7 | 1,98 | 0,85 |
| 90 | ChaB | YP_006908598.1 | 75 | 45,33 | 2 | 2,98 | 1,28 |
| 92 | **VP39** | YP_006908600.1 | 293 | 75,43 | 21 | 232,27 | 100,00 |
| 93 | **ODV-EC27** | YP_006908601.1 | 284 | 24,65 | 8 | 3,39 | 1,46 |
| 94 | BRO | YP_006908602.1 | 359 | 5,85 | 2 | 0,23 | 0,10 |
| 95 | Epap95 | YP_006908603.1 | 73 | 69,86 | 5 | 176,83 | 76,03 |
| 96 | **PIF8** | YP_006908604.1 | 567 | 14,10 | 7 | 1,22 | 0,52 |
| 98 | **Ac81** | YP_006908606.1 | 191 | 23,04 | 5 | 1,61 | 0,69 |
| 99 | **GP41** | YP_006908607.1 | 286 | 74,82 | 23 | 87,59 | 37,66 |
| 100 | **Ac78** | YP_006908608.1 | 88 | 44,32 | 2 | 16,78 | 7,22 |
| 101 | **VLF-1** | YP_006908609.1 | 368 | 35,87 | 11 | 3,06 | 1,32 |
| 103 | Ac75 | YP_006908611.1 | 149 | 56,38 | 9 | 7,11 | 3,06 |
| 106 | **DNA Pol** | YP_006908614.1 | 1068 | 2,72 | 1 | 0 | 0,00 |
| 109 | **PIF6** | YP_006908617.1 | 148 | 41,22 | 5 | 2,59 | 1,11 |
| 113 | FP25K | YP_006908621.1 | 146 | 19,86 | 3 | 0,23 | 0,10 |
| 115 | DNA Lig | YP_006908623.1 | 534 | 4,87 | 1 | 0,08 | 0,03 |
| 123 | Epap123 | YP_006908630.1 | 102 | 6,87 | 1 | 0,47 | 0,20 |
| 126 | Epap126 | YP_006908633.1 | 347 | 45,82 | 11 | 3,49 | 1,50 |
| 127 | Epap127 | YP_006908634.1 | 69 | 30,43 | 2 | 1,15 | 0,49 |
| 129 | **VP1054** | YP_006908636.1 | 344 | 11,05 | 2 | 0,1 | 0,04 |
| 133 | ME53 | YP_006908640.1 | 373 | 9,91 | 2 | 0,22 | 0,09 |
| - | **Ac110** | This study | 47 | 29.79 | 1 | - | - |

% Coverage: percentage of the protein sequence covered by identified peptides.
emPAI: exponentially modified Protein Abundance Index.
ORF: Open Reading Frame, as numbered in EpapGV genome map published by Ferrelli *et al* [7].

In addition to identifying the components of the OBs, we estimated the relative abundance of each protein; to this end we calculated the emPAI value proposed by Ishihama *et al* [26] for each protein. The emPAI value for the major capsid protein VP39 was used to normalize protein abundance (Table 1). Taking a cutoff value of at least 10 % VP39 emPAI, the most abundant proteins are GP41, Granulin, ODV-E18, SOD and Epap95, together with VP39. The major capsid component VP39, the tegument protein GP41 and the major component of OB matrix granulin were expected to be among the most abundant proteins due to their known structural function. ODV-E18 ortholog in AcMNPV is an essential protein for BV production that also localizes to the ODV membrane [27]. Cu-Zn superoxide dismutase (SOD) activity in virion preparations of Chlorovirus PBCV-1 has been recently associated with reactive oxygen species reduction during the early stages of virus infection [28]. Finally, Epap95 a protein with orthologs in all the members of the genus *Betabaculovirus*, has been consistently detected in the granuloviruses of *Clostera anachoreta* (ClanGV) and *Pieris rapae* (PiraGV) as a component within ODVs.

Betabaculovirus gene content remains poorly characterized, with a large number of hypothetical genes predicted by phylogenomics methods. Some of these genes lack *bona fide* experimental evidence to confirm the actual existence of their putative protein products. Our proteomic data confirmed the presence of translation products for 10 hypothetical proteins annotated in the genome of EpapGV, namely, Epap10, Epap40, Epap48, Epap49, Epap62, Epap71, Epap95, Epap123, Epap126 and Epap127.

### Short peptides encoded in EpapGV genome that do not belong to annotated ORFs

To identify possible unannotated proteins, we searched our spectral data against a theoretical database comprising all translation products predicted in the six reading frames of EpapGV genome sequence (we included all possible ORFs, without introducing a minimal size criterion). We detected seven peptides that mapped to the EpapGV genome but did not belong to the set of annotated ORFs [7]. Their sequence and genomic location are detailed in S1 Appendix. One of these peptides matches an unannotated 47 amino acids long ORF overlapping *epap51* but in the opposite orientation. We further examined the presence of this novel ORF in other members of the family *Baculoviridae* and found that it is an ortholog of the core gene *ac110* [29]. This gene has been described as the *per os* infectivity factor 7 (*pif7*) and its product has only been detected in the proteome of HearNPV ODV [18] and EpapGV (this study). Genomic localization and orientation of this *ac110*-like gene is conserved within *Betabaculovirus*, providing additional evidence about its evolutionary conservation.

The remaining six peptides overlap with annotated ORFs (*chitinase*, *dna ligase* and *granulin*) or intergenic regions. Two peptides were found between ORFs *epap48* and *epap49* and one peptide between *epap61* and *epap62.* TBLASTN was used to find putative homologous unannotated peptides in other baculovirus genomes. Only the peptides overlapping with *chitinase* and *granulin* are conserved in homologous *loci* in GV and NPV genomes (S1 Appendix).

Remarkably, peptides between *epap48* and *epap49* almost cover the entire 145 bp intergenic sequence (Fig 2). *Epap48* encodes a 446 amino acid long protein that is conserved in the betabaculoviruses. The putative translation product of *epap49* is a large protein composed of 1465 residues with no orthologs detected in other baculoviruses. Mapped peptides are located in the same reading frame as the translation product of *epap49*, but no methionine codon has been found in frame (Fig 2). One hypothesis that could explain the presence of these peptides is that Epap48 and Epap49 may be expressed as a fusion protein due to a putative +1 frameshifting event near the C-terminus of Epap48; further experimental validation of this potential fusion protein will be needed to confirm this hypothesis.

**Fig. 2.**
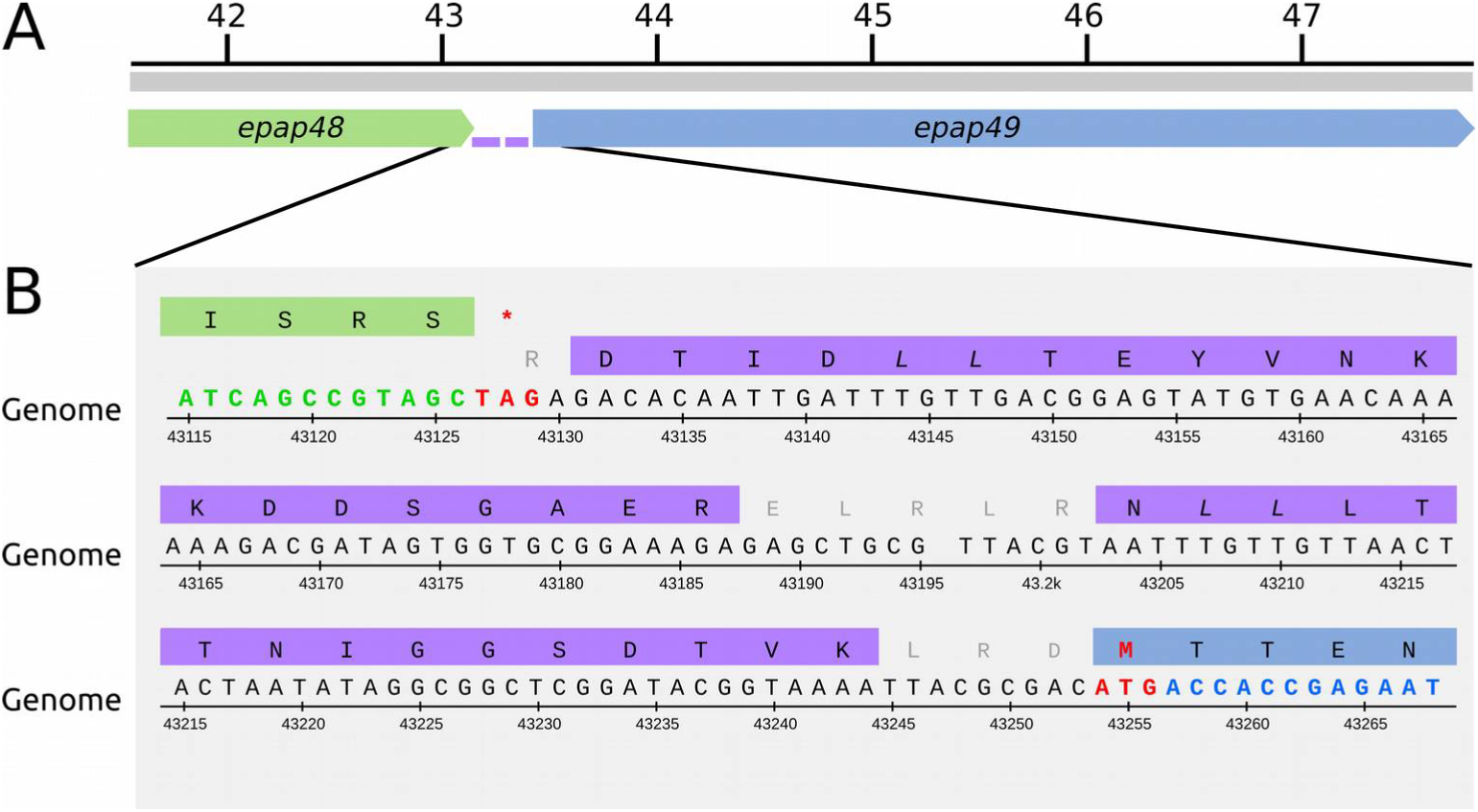
Putative fusion protein Epap48-Epap49. (A) Genomic locus containing *epap48* (green) and *epap49* (blue) genes. The two peptides detected by MS inside the intergenic region are depicted in purple. (B) Genome sequence and translation frame for each gene. Start and stop codons are shown in red (nucleotide numbers are those from NCBI accession number NC_018875).

### Conservation of structural proteins in the family Baculoviridae

The reports of ODV proteomes belonging to several baculoviruses were used to evaluate the conservation of the viral particle composition in this family. To date, eight proteomic studies were carried out on ODV, including five members of the genus *Alphabaculovirus* (AcMNPV, AgMNPV, ChchNPV, MabrNPV and HearNPV), two of the genus *Betabaculovirus* (ClanGV and PiraGV) and one of *Deltabaculovirus* (CuniNPV). Our study expands this data with the proteins present in the EpapGV OBs. Sequences of proteins detected in occluded virions of baculovirus were used to construct groups of orthologous proteins (S1 Table). For each of these orthogroups we scored the number of proteomes in which they are present as a measure of their conservation. We assigned them a class based in the phylogenetic conservation of their coding sequence (core, lepidopteran-specific, genus-specific and virus specific) (Fig 3A). Most conserved protein groups (present in a larger number of proteomes) are enriched in core and lepidopteran-specific gene products. In contrast, proteins specific to a small set of proteomes are encoded by genus-specific and virus-specific genes.

**Fig. 3.**
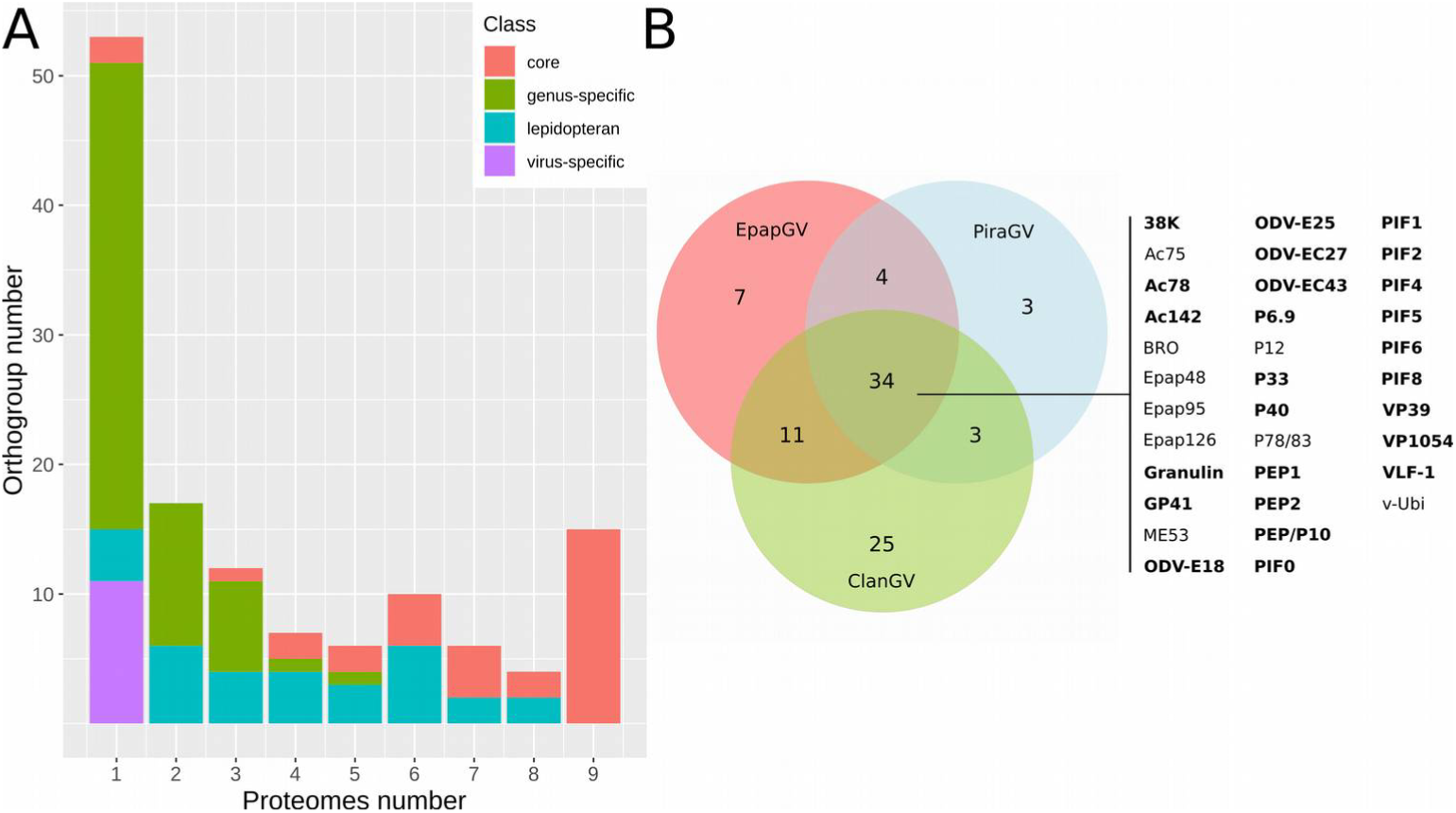
Baculovirus proteome comparison. (A) Protein conservation in baculovirus occluded virion proteomes. Protein orthogroups derived from proteomics datasets were scored according to the number of proteomes in which they were detected. Gene class distribution (core genes, lepidopteran baculovirus-specific, genus-specific, specie-specific) of orthologs groups within *Baculoviridae* is highlighted in different colours. (B) Proteins detected in proteomes of betabaculoviruses were grouped in sets of orthologs and represented using a Venn diagram. A set of 34 proteins is present in all three viruses [20, 21, this study]; core gene products are highlighted in bold. Two of these protein clusters being specific of *Betabaculovirus* (Epap48 and Epap95).

The betabaculovirus proteomes (EpapGV, ClanGV and PiraGV) were compared using a Venn diagram (Fig 3B). From the proteins present in all three viruses, BRO, Epap48, Epap95 and Epap126 are the only orthogroups without functional characterization. Interestingly, Epap95 is one of the most abundant proteins according to emPAI values. Additionally, Epap126 is shared between group II alphabaculoviruses and betabaculoviruses (except for ClanGV) (S1 Table). On the other hand, Epap10, Epap49 and Epap62 are proteins unique to EpapGV OBs. Remarkably, *epap10* orthologs are encoded only in five alphabaculoviruses that infect insects of the family *Tortricidae*, *Choristoneura fumiferana* NPV, *Choristoneura occidentalis* NPV, *Choristoneura rosaceana* NPV, *Cryptophlebia peltastica* NPV and *Epiphyas postvittana* NPV. This could be the product of an ancestral horizontal gene transfer between alphabaculoviruses and betabaculoviruses that coinfected the same host, based on gene conservation evidence.

## Discussion

Occluded virions are responsible for baculovirus primary infection. Proteome of OBs is related with oral infectivity, providing relevant information about conserved components potentially associated with midgut infection. Until now, the proteomes of ClanGV and PiraGV ODVs have been interrogated using MS-based techniques [20, 21]. These viral species are phylogenetically distant to EpapGV [7]. We explored possible divergence in protein composition employing a bottom-up proteomic approach to characterize the protein content of EpapGV OBs. A diagram of the EpapGV virion particle summarizing qualitative and semi-quantitative composition is shown in Fig 4. Virion components can be grouped in five classes based in their localization: 18 nucleocapsid proteins, 15 ODV envelope proteins, 5 occlusion matrix proteins, 1 tegument protein and 17 proteins of undefined localization. Comparisons across virion proteomes available for members of the family *Baculoviridae* highlighted the conservation of several structural components forming the mature virion. On the other hand, comparative genomics highlight the conservation of a collinear genomic region for lepidopteran-infecting baculoviruses [30]. Combined genomics and proteomics information suggests that this *locus*, compared to the rest of genome, is densely populated by protein coding sequences corresponding predominantly to structural polypeptides (Fig 1 and S1 Fig). Intriguingly, the product of the core gene desmoplakin could not be detected in the betabaculovirus structural proteomes but it has been reported for alphabaculoviruses; we do not know if this is related to a different localization of this protein within betabaculovirus or due to technical reasons. In AcMNPV, desmoplakin has been implied in the segregation of nucleocapsids destined to build BV (which are ubiquitinated) and ODV (non ubiquitinated) [31].

**Fig. 4.**
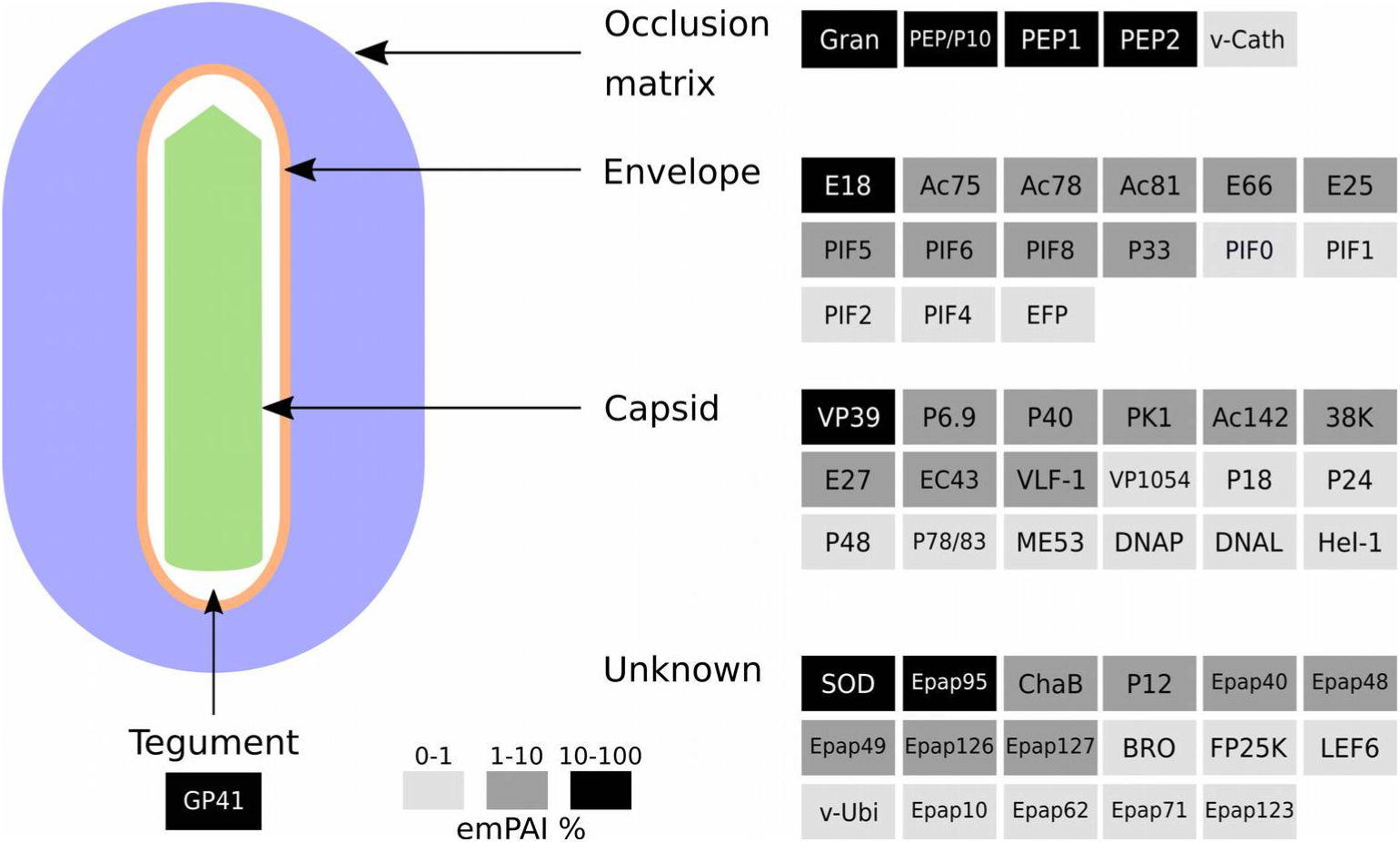
Schematic model of EpapGV OB. Qualitative and semi-quantitative virion model based on our proteomic data and localization described in published reports. Protein levels were estimated using the emPAI value and expressed as relative value with respect to the major capsid protein VP39.

It has been previously reported that proteins related to viral DNA metabolism and DNA binding capacity may be retained in the virion [15]. In the present study we could identify the DNA polymerase, DNA ligase and Helicase-1 in OBs. For other baculoviruses IE1, Alk-Exo, LEF1 and LEF3 have been detected also. This reinforces the idea that the viral DNA is associated with various proteins (in addition to the major condensing protein P6.9) inside the viral capsid.

The envelope that surrounds the ODV morphotype is especially adapted for primary infection of the insect midgut and presents a complex complement of proteins. These can be classified in two functional groups, those required for virion envelopment and those related with oral infectivity. Envelope morphogenesis begins with the formation of intranuclear microvesicles (IMV) derived from the inner nuclear membrane and the association with viral capsids. The ODV membrane proteins Ac75 and P18 are necessary for the generation of these IMV [32, 33]. Subsequently, envelopment of assembled nucleocapsids requires the ODV proteins Ac78, Ac81, Ac142, ODV-E25, ODV-EC43, P33 and P48 to form mature OBs [34-40]. On the other hand, several ODV membrane proteins are members of the PIF complex (PIF0, PIF1, PIF2, PIF3, PIF4, PIF5, PIF6 and PIF8); this molecular complex is the main effector of oral infection in the insect midgut. These proteins are encoded by core genes conserved in all the members of the *Baculoviridae* family [41].

The biological relevance of the betabaculovirus-specific orthogroups Epap48 and Epap95 within OBs is currently unknown. Moreover, the high content of Epap95 in the OBs may also be biologically relevant. On the other hand, Epap126 orthogroup is present in group II alphabaculoviruses and betabaculoviruses, which represents a conserved protein potentially involved in oral infection.

Baculovirus genomes are densely populated with coding sequences (overlapping in several cases) and contain short intergenic regions [2]. A recent study has described the transcriptional landscape of baculovirus infection, demonstrating the existence of several polycistronic and overlapping viral transcripts [42]. Together with other technologies, proteogenomic mapping is a valuable tool to improve the annotation of these complex coding regions. This approach has been used in proteome research for several virus families [43], but was applied only for one baculovirus, AgMNPV [44]. We identified seven peptides that do not map to previously annotated coding regions. One of these peptides turned to be an ortholog of *ac110*; this ORFs overlapped with the coding sequence of *epap51.* Moreover, the presence of peptides derived from alternative frames inside the coding regions of *granulin* and *chitinase* raises the question about the underlying complexity of baculovirus transcription and translation processes.

Surprisingly, two unmapped peptides were found to be encoded in the intergenic region of *epap48* and *epap49* and suggest the presence of a putative fusion product between these proteins. Epap49 is the largest protein annotated in EpapGV genome; it is 1465 amino acids long. As previously reported, it was difficult to annotate this as a hypothetical protein due to its atypically large size, absence of homologous proteins in Genbank and lack of known promoter motifs [7]. Also, it was noted that large proteins were coded in similar locations in the genomes of ChocGV and HearGV, 1144 and 1279 amino acids long, respectively [45, 46]. In the case of ChocGV it was not annotated in the genome [45] and in HearGV it was found to be a fusion of ORFs homologous to XecnGV 47 and 48 [46]. In this study, we found evidence that Epap49 is actually translated.

## Conclusion

The protein composition of EpapGV OBs was interrogated using an MS-based proteomic approach. A total of 56 proteins have been detected in the EpapGV occluded virion, suggesting the presence of a highly conserved protein profile in baculoviral OBs. We identified Epap95, a betabaculovirs-specific protein, as a highly abundant capsid component. This protein represents an interesting candidate for further functional studies to explore its role in betabaculovirus pathogenesis. Through proteogenomic search we could detect a non-annotated coding region with a high degree of sequence identity to *ac110.* In addition, our data strongly suggest the translation of a putative fusion protein involving Epap48 and Epap49. Our study highlight the usefulness of MS proteomics to characterize the protein complement of the viral particle and the possibility to improve genome annotation through experimental evidence for translation of predicted coding regions.

## Acknowledgements

The authors thank Dr. Silvia Margarita Moreno and Dr. María Pía Valacco, from the Centro de Estudios Químicos y Biológicos por Espectrometría de Masas (CEQUIBIEM-CONICET-FCEN-UBA), for their help with sample preparation protocols, data acquisition and subsequent analysis. This work was supported by grants from the Agencia Nacional de Promoción Científica y Tecnológica (ANPCyT) and UNLP to Víctor Romanowski.

## Supplementary information

**S1 Appendix.** EpapGV unannotated peptides detected by MS.

**S1 Figure.** Parity plot of sequences coding for structural proteins present in AcMNPV, ChchNPV and PiraGV against EpapGV.

**S1 Table.** Proteomic profiles of baculovirus occluded virions proteomes.

